# Genome-Wide Association Studies Identify 15 Genetic Markers Associated with Marmite Taste Preference

**DOI:** 10.1101/185629

**Authors:** Thomas R. Roos, Nikolay A. Kulemin, Ildus I. Ahmetov, Avi Lasarow, Keith Grimaldi

## Abstract

Marmite is a popular food eaten around the world, to which individuals have commonly considered themselves either “lovers” or “haters”. We aimed to determine whether this food preference has a genetic basis.Weperformed a genome-wide association study (GWAS) for Marmite taste preference using genotype and questionnaire data froma cohort of 261 healthy adults. We found 1 single nucleotide polymorphism (SNP) associated with Marmite taste preferencethat reached genome-wide significance (p<5x10-^8^) in our GWAS analyses. We found another 4 SNPsassociated with Marmite taste preference that reached genome-wide significance (p<5x10-^8^) in at leastoneGWAS and/or for at least one phenotype analysed. Moreover, we identified 10 additional SNPs potentially associated with Marmite taste preference through candidate gene analysis. Our results indicate that there is a genetic basis to Marmite taste preference and we have identified 15 genetic markers for this trait. Overall, we conclude that Marmite taste preference is a complex human trait influenced by multiple genetic markers, as well as the environment.

**Summary of Main Results:** 1. Marmite taste preference is a complex human trait with many factors influencing whether an individual loves or hates Marmite.
2. The relative contribution of genetics versus environment (ie. heritability) for Marmite taste preference is unknown.
3. The genetic contribution to Marmite taste preference involves multiple genetic markers each contributing a small amount (ie. the trait is polygenic). There is not one single Marmite gene with a large contribution like in the case of the *TAS2R38*gene and bitter taste perception.
4. We have found a total of 15 SNPs associated with Marmite taste preference: 5 SNPs by a genetic-association screen atgenome-wide significance, and 10 SNPs by a candidate gene approach at nominal significance.
5. We did not find an association between the *TAS2R38*bitter taste receptor gene and Marmite taste preference.
6. It is important to independently replicate the findings of this study in order to validate these genetic markers and get a more accurate idea of their true effect on Marmite taste preference.

## Introduction

There is evidence in the nutrigenomics literature showing that there is a genetic basis for human taste perception. In particular that single nucleotide polymorphisms (SNPs) in taste receptor genes are associated with bitter tasting ability [Reed et al, Physiol Behav, 2006] and umami tasting ability [Chen et al, Am J Clin Nutr, 2009].

For example, the genetic contribution to bitter taste perception is estimated as 55-85% from family and twin studies [Hansen et al, Chem Sense, 2006]. The remaining contribution comes from demographic factors (ie. gender, age, and ethnicity) and environmental factors (ie. diet, saliva composition, tongue morphology, and smoking) [Mennella et al,BMC Genet, 2010]. The Taste Receptor 2 Member 38 (TAS2R38) is a well-studied G-protein coupled taste receptor that is linked to bitter taste perception in humans. It is encoded by the *TAS2R38*gene, which has 3 SNPs (Pro49Ala, Ala262Val, Val296Ile) associated with bitter tasting ability [Bachmanov et al, Annu Rev Nutr, 2007]. These 3 SNPs create 2 common haplotypes: 1) PAV, the dominant taster group, and 2) AVI, the recessive non-taster group. Individuals with 1 or 2 copies of the PAV haplotype are more sensitive to bitter taste, while individuals with 2 copies of the AVI haplotype are less sensitive. While there are over 25 taste receptors thought to contribute to bitter taste perception [Tepper et al, Nutrients, 2014], the *TAS2R38*gene accounts for largest genetic contribution to the trait [Kim et al,Science, 2003].

It is commonly known that some people “Love” Marmite while others “Hate” the taste, but the reasons for this are unknown. Some individuals consider Marmite to taste bitter, others consider it to taste savory, and yet others consider it to taste neutral. This inter-individual difference in taste perception could contribute to whether people describe themselves as Marmite “Lovers” or “Haters” or could influence how people rate the taste of Marmite. The relative contributions of genetics and the environment to this trait are unknown.

The primary aim of this study was to investigate whether genetics contributes to taste perception of Marmite and/or taste preference of Marmite. The secondary aim of this study was to determine whether genetic markers already known to be associated with taste and/or olfactory function are also associated with liking or disliking Marmite. Importantly, this is the first study to show that genetic variations are associated with Marmite taste preference.

## Results

### Genome-Wide Association Studies for Marmite Taste Preference

Genome-wide association studies were performed to identify SNPs associated with Marmite taste preference in an additive genetic model (Methods). Our cohort consisted of 261 total study subjects, with a subgroup of 213 subjects of white European descent [Table 1]. We tested 2 phenotypes related to the Marmite taste preference trait for genetic associations using basic association models and logistic/linear regression models. We controlled for sex, age, and smoking status as covariates in the model, as well as adjusting results for genomic inflation. For each of the ~650,000 directly genotyped SNPs, we obtained p-values and odds ratios and/or beta coefficients from the models. To assess significance, we set our statistical thresholds in accordance with standard practice: genome-wide significance (p<5x10-^8^), Bonferroni corrected significance (p<7.78x10^8^), and nominal significance (p<1x105). To assess effect size, we set ranges in accordance with standard practice for odds ratios (OR): small effect (OR=1.0-1.5), medium effect (OR=1.5-6.5), and large effect (OR>6.5).

**Table 1.** Demographic factors of the study population used in genome-wide association analyses of Marmite taste preference.

| **COHORT** | **ALL** | **EUR** |
| --- | --- | --- |
| **Subjects** [N (%)] | 261 (100%) | 213 (100%) |
| **Gender/Sex<sup>a</sup>** |  |  |
| Female | 136 (52.1%) | 116 (54.5%) |
| Male | 125 (47.9%) | 97 (45.5%) |
| Transgender/Other | 0 (0.0%) | 0 (0.0%) |
| **Age<sup>b</sup>** |  |  |
| Overall [$\mu$ ($\sigma$)] | 28.8 (7.5) | 28.9 (7.8) |
| 18-24 | 87 (33.3%) | 73 (34.3%) |
| 25-34 | 130 (49.8%) | 104 (48.8%) |
| 35-44 | 32 (12.3%) | 24 (11.3%) |
| 45-54 | 9 (3.4%) | 9 (4.2%) |
| 55-64 | 3 (1.2%) | 3 (1.4%) |
| >65 | 0 (0.0%) | 0 (0.0%) |
| **Ethnicity/Race<sup>c</sup>** |  |  |
| White European | 213 (81.6%) | 213 (100%) |
| Mixed Ethnicity | 20 (7.7%) | N/A |
| Asian | 16 (6.1%) | N/A |
| Black/African/Caribbean | 9 (3.4%) | N/A |
| Other | 3 (1.2%) | N/A |
| **Nationality<sup>d</sup>** |  |  |
| English/British | 192 (73.5%) | 153 (71.8%) |
| Other United Kingdom | 23 (8.8%) | 23 (10.8%) |
| Other European | 26 (10.0%) | 24 (11.3%) |
| Other | 20 (7.7%) | 13 (6.1%) |
| **Smoking<sup>e</sup>** |  |  |
| Never | 147 (56.3%) | 118 (55.4%) |
| Past | 60 (23.0%) | 51 (23.9%) |
| Current | 54 (20.7%) | 44 (20.7%) |

<sup>a</sup> Gender/Sex as determined by self-reported gender and confirmed by subjects genetic data.
<sup>b</sup> Age at subject enrollment in the Marmite Genetics study cohort, reported as mean age with standard deviation and broken into standardized age groups.
<sup>c</sup> Ethnicity/Race as determined by self-reported ethnicity according to UK NHS Census standardized categories.
<sup>d</sup> Nationality as determined by self-reported nationality according to UK NHS Census standardized categories or subjects free response answer, then collapsed to the 4 categories presented.
<sup>e</sup> Smoking as determined by self-reported smoking status.

Our results identified 1 SNP (rs11122562) that reached genome-wide significance (p<5x10-8) in multiple analyses [Table 2]. We found that this SNP is significantly associated with whether people self-describe themselves as a Marmite lover, hater, or neutral, and likely has a moderate to large effect size. In a case/control model (lovers vs haters, either including neutrals with lovers or excluding neutrals), our basic association results (p=6.84x10^−8^, OR=8.26) and regression model results (p=2.86x10^−6^, OR=6.92) indicate a significant association.In a continuous model (lovers vs neutral vs haters), our basic association results (p=1.50x10-8) and regression model results (p=6.65x108) also both indicate a significant association. We found that this SNP is associated with how people rate the taste of Marmite on a five point Likert scale (love, like, neutral, dislike, hate). In a continuous model (using all 5 categories), our basic association results (p=4.45x10-9) and regression model results (p=3.04x109) both reach genome-wide significance. In a continuous model (using 3 categories, grouping love/like and dislike/hate), our basic association results (p=1.78x10-8) and regression model results (p=4.98x109) also both reach genome-wide significance. Moreover, both of these associations held true when including the entire cohort or just the European subgroup of the cohort. Overall, these results show that rs11122562 is significantly associated with both phenotypes of the Marmite preference trait.

**Table 2.** Top SNPs associated with Marmite taste preference from genome-wide association analyses.

| **SNP** | **Gene** | **Phenotype** | **P-Value** |
| --- | --- | --- | --- |
| rs11122562 | COG2 | Love Hate | 1.50E-08 |
| rs11122562 | COG2 | Rate Marmite | 4.45E-09 |
| rs79342650 | IRF1/IL5 | Rate Marmite | 1.79E-08 |
| rs10758801 | GLDC | Rate Marmite | 5.44E-08 |
| rs17335506 | CEP290 | Love Hate | 2.53E-08 |
| rs75189446 | ITNS1/RUNX1 | Rate Marmite | 4.89E-08 |

Our results identified 4 additional SNPs (rs79342650, rs75189446, rs10758801, rs17335506) that reached multiple hypothesis corrected significance (p<7.78x10-8) through basic association modeling for at least one phenotype [Table 2]. Notably, these 4 SNPs also reached nominal significance (p<1x105) through regression modeling, after accounting for covariates, and after adjusting for genomic inflation. The first 2 SNPs are associated with how people rate the taste of Marmite in the full cohort: rs79342650 (p=1.79x10-^8^) and rs75189446 (p=4.89x10-^8^). The third SNP is associated with how people rate the taste of Marmite in the European subgroup: rs10758801 (p=5.44x10-8). The fourth SNP is associated with Marmite “lover” vs “hater” status in the full cohort: rs17335506 (p=2.53x10^−8^). Overall, these results show that there is most likely multiple SNPs whose genetic effects contribute to the Marmite taste preference trait.

We investigated these 5 SNPs for evidence that they may affect the expression or coding capacity of neighboring genes, consistent with being variants that are causal for affecting Marmite taste preference. None of the SNPs are found in exons of genes or are annotated as directly affecting protein-coding regions of the genome. However, 3 of the SNPs are in intronic regions of the genome. Specifically, rs10758801 (*GLDC*),rs17335506 (*CEP290*),and rs75189446 (*ITNS1/RUNX1).*These genes are not known to be involved in biological pathways related to taste or olfactory sensation. None of the SNPs are annotated as acting as expression quantitative trait loci (eQTLs). These data suggest that none of the 5 SNPs has strong evidence for being a causal mutation.

Next we investigated whether these 5 SNPs might play a regulatory role in gene expression by using data from the ENCODE project. In general, when transcription factors bind to DNA at the promoter region of genes, they are associated with distinct patterns of DNA methylation. Moreover, enhancer sites located near genes may also aid in transcription factor binding. Thus, SNPs located in these regions may consequently affect gene expression. One SNP in our study, rs79342650, is located in the promoter region of its adjacent gene (*IRF1*)and in an H3K4me peak. Another SNP in our study, rs10758801, is located in a predicted enhancer region, possibly for its adjacent gene (*GLDC*),and in an H3K4me peak. These data suggest that these two SNPs are located in regulatory regions and might have an effect on expression of nearby genes.

### Genome-Wide Association Studies for Bitter Taste Ability

Genome-wide association studies were performed to identify SNPs associated with bitter tasting ability in an additive genetic model (Methods). As above, we tested 2 phenotypes related to the ability to taste bitter compounds (ie.PTC) for genetic associations using basic association models and logistic/linear regression models.

Our results identified 3 SNPs (rs713598, rs1726866, rs10246939) that reached genome-wide significance (p<5x10-8) in all analyses [Table 3]. The first SNP, rs713598, showed strong association and a medium effect size for both phenotypes of whether PTC tasted bitter (p=4.08x10^−18^, OR=5.17) and rating how PTC tasted (p=4.54x10^−16^, OR=5.04). The second SNP, rs1726866, also showed strong association and a medium effect size for both phenotypes of whether PTC tasted bitter (p=2.52x10-17, OR=4.83) and rating how PTC tasted (p=4.45x10^−17^, OR=4.66).And the third SNP, rs10246939, also showed strong association and a medium effect size for both phenotypes of whether PTC tasted bitter (p=4.85x10^−18^, OR=5.05) and rating how PTC tasted (p=6.13x1017, OR=4.71). These associations held true in both basic and regression models, and in both the entire cohort and the European subgroup. Overall, these results show that these 3 SNPs are strongly associated with bitter tasting ability.

**Table 3.** Top SNPs associated with bitter tasting ability from genome-wide association analyses.

| **SNP** | **Gene** | **Phenotype** | **P-Value** | **Odds Ratio** |
| --- | --- | --- | --- | --- |
| rs713598 | TAS2R38 | PTC Bitter | 4.08E-18 | 5.17 |
| rs713598 | TAS2R38 | Rate PTC | 4.54E-16 | 5.04 |
| rs1726866 | TAS2R38 | PTC Bitter | 2.52E-17 | 4.83 |
| rs1726866 | TAS2R38 | Rate PTC | 4.45E-17 | 4.66 |
| rs10246939 | TAS2R38 | PTC Bitter | 4.85E-18 | 5.05 |
| rs10246939 | TAS2R38 | Rate PTC | 6.13E-17 | 4.71 |

All 3 of these SNPs are located on chromosome 7 in the *TAS2R38*gene and are the same 3 SNPs that have been previously associated with the ability to taste bitter compounds such as PTC. Our results confirm previous findings that genetics strongly affects the bitter taste trait and whether people are able to taste bitter compounds in foods. Importantly, this result serves as a positive control to indicate that we did not have severe bias in our cohort, in the data we collected, or in our analysis methodology.

### Candidate Gene Studies for Marmite Taste Preference

In addition to our hypothesis-free GWAS approach, we conducted candidate gene analyses using hypotheses based upon previous biological and genetic knowledge. We created a list of 252 genes involved in taste and/or olfactory sensation. This list included both protein-coding genes and pseudogenes annotated as related to the biological processes of: taste(n=60), taste transduction (n=69), odor sensing (n=70), and hormone signaling pathways (n=53). Nearly all these genes code for taste receptors, olfactory receptors, and neurotransmitter receptors found in the mouth, nose, gut, and brain of humans.

We used the genetic data from our study to evaluate this set of candidate genes for association with Marmite taste preference. We performed genetic association by comparing allele frequencies in Marmite “lovers” to Marmite “haters” for the ~650,000 genotyped SNPs in a case/control model. For each SNP that reached nominal significance (p<0.05) by chi-squared and/or Fishers exact analysis, we annotated the gene that the SNP was located in (or the closest gene nearby in the genome). Next we cross-referenced the two lists to select for genes from the list of 252 candidates that included SNPs with nominally significant p-values. Ultimately we identified 10 candidate genetic markers: rs779710 in *GRM7*(p=0.0007); rs113693459 in *ASIC2*(p=0.0011); rs10243170 *in ADCY1*(p=0.0011); rs857940 in *OR6K3*(p=0.0056); rs17214874 in *MAOA*(p=0.0067); rs10256873 in *GRM8*(p=0.0070); rs2854038 in *CCKAR*(p=0.0145); rs4657718 *in ADCY10*(p=0.0149); rs4906728 in *UBE3A*(p=0.0167); and rs28610409 in *OR2T10*(p=0.0293) [Table 4].

**Table 4.** Top SNPs associated with Marmite taste preference from candidate gene analyses.

| **SNP** | **Gene** | **P-Value** |
| --- | --- | --- |
| rs779710 | GRM7 | 0.0007 |
| rs113693459 | ASIC2 | 0.0011 |
| rs10243170 | ADCY1 | 0.0047 |
| rs857940 | OR6K2/OR6K3 | 0.0056 |
| rs17214874 | MAOA | 0.0067 |
| rs10256873 | GRM8 | 0.0070 |
| rs2854038 | CCKAR | 0.0145 |
| rs4657718 | ADCY10 | 0.0149 |
| rs4906728 | UBE3A | 0.0167 |
| rs28610409 | OR2T10 | 0.0293 |

Furthermore, we used the genetic data from our study data to evaluate the *TAS2R38*gene for association with Marmite taste preference. Our hypothesis was that people who have the ability to taste bitter compounds (especially “supertasters” who have a strong reaction to PTC) are more likely to dislike the taste of Marmite. When tested independently, none of the 3 individual SNPs in *TAS2R38*(rs713598, rs1726866, rs10246939) were significantly associated with either Marmite taste preference phenotype in either the full cohort or the European subgroup [Table 5]. We found no association for the 3 SNPs (p=0.45, p=0.40, p=0.48) respectively, with Marmite “lover” vs “hater” status. Likewise, we found no association for the 3 SNPs (p=0.42, p=0.28, p=0.32) respectively, with rating the taste of Marmite. Next, we tested the PAV haplotypes of the *TAS2R38*gene combining the genotypes at these 3 SNPs in both additive and dominant genotypic models. Again we found no significant association with Marmite “lover” vs “hater” status (analysed as a 3 or 2 level phenotype respectively), using an additive model (p=0.36 or p=0.86) or a dominant model (p=0.42 or p=0.85). Thus, the data do not show a correlation between bitter tasting ability and Marmite taste preference.

**Table 5.** TAS2R38 candidate SNPs association with Marmite taste preference from candidate gene analyses.

| **SNP/Haplotype** | **Gene** | **Phenotype** | **P-Value** |
| --- | --- | --- | --- |
| rs713598 | TAS2R38 | Love Hate | 0.45 |
| rs713598 | TAS2R38 | Rate Marmite | 0.42 |
| rs1726866 | TAS2R38 | Love Hate | 0.40 |
| rs1726866 | TAS2R38 | Rate Marmite | 0.28 |
| rs10246939 | TAS2R38 | Love Hate | 0.48 |
| rs10246939 | TAS2R38 | Rate Marmite | 0.32 |
| PAV (additive) | TAS2R38 | Love Hate (3) | 0.36 |
| PAV (additive) | TAS2R38 | Love Hate (2) | 0.86 |
| PAV/* (genotypic) | TAS2R38 | Love Hate (3) | 0.42 |
| PAV/* (genotypic) | TAS2R38 | Love Hate (2) | 0.85 |

## Discussion

With the advent of large-scale genotyping programs, it is now possible to screen the entire genome for genetic variants associated with tasting ability and food preference. Furthermore, because the genotype data includes most of the common polymorphisms that are known, a genome-wide screen reports the strongest associations in the genome in an unbiased manner. Here, we have performed a study to find DNA single nucleotide polymorphisms (SNPs) associated with Marmite taste preference using a cohort of 261 individuals. To date, this is the first gene association study for Marmite nutrigenomics. Our data provide new insights regarding the differences between individuals who are Marmite “Lovers” and those who are Marmite “Haters” and show that there is a genetic basis to Marmite taste preference.

Notably, the top SNP from our study (rs11122562) reached genome-wide significance (p<5x10^−8^) for association with Marmite taste preference. This association is valid for both the lover vs hater and rating the taste of Marmite phenotypes, for the entire cohort and the European subgroup, and after including covariates and adjustments. Additionally, we found 4 SNPs that reached multiple hypothesis corrected significance (p<7.78x10^−8^) in one or more analyses. It is unclear if any of these are causal SNPs that directly affect Marmite taste preference or if they are simply linked SNPs (ie. passive bystanders). In our study, these 5 SNPs have moderate to large effect sizes (OR=5.0-20.0) on the Marmite taste preference trait. Due to the small sample size of our study these effect estimates are most likely inflated from the true effect.

An assessment of previous genetic association studies found that 73% of SNP associations with borderline significance (between p≤1x10^−7^ and p>5x10^−8^) were replicated in subsequent studies [Panagiotou et al, Int J Epidemiol, 2012]. For SNP associations with suggestive significance (between p≤1x10^−6^ and p>1x10^−7^) the replication rate is likely to be much smaller, but not negligible. Thus, it is important to note that the SNPs found in this study will need to replicated to validate their association with Marmite preference traits. Moreover, it is likely that there are many common SNPs with small effects (OR<1.5) that are associated with Marmite preference, but which were not detected in our study. To identify SNPs with small effect sizes in a genome-wide screen, it will be necessary to increase the sample size or meta-analyze these results with results from a replication study [Panagiotou et al, Annu Rev Genomics Hum Genet, 2013].

We identified 10 candidate genetic markers associated with Marmite taste preference phenotypes that reached nominal significance (p<0.05). These SNPs and associated genes have relevance to taste and odor reception pathways, thus might play a biological role in Marmite preference. Evidence from many other genetic association studies suggests that candidate gene associations need to be independently replicated, otherwise their credibility is low [Siontis et al, EJHG, 2010].

An important finding from our study is that we did not find any association between the *TAS2R38*candidate gene (either the 3 SNPs individually or the PAV haplotype) and Marmite taste preference. Our study validated previous studies in showing that these SNPs have a strong association (p<1x10^−15^) and moderate effect size (OR=4.5-5.5) on an individuals bitter tasting ability, as tested by PTC. If bitter tasting ability has a moderate to large effect on Marmite taste preference (OR>1.5), power calculations indicate that we would have had an 80-85% chance to find common SNPs with true genetic associations to this trait [Ioannidis et al, Epidemiology, 2011]. Thus, our data do not support the hypothesis that the *TAS2R38*gene or bitter tasting ability plays a significant role in whether people love or hate Marmite. Future investigation into other taste receptor genes and olfactory receptor genes is warranted to learn more about the genetic basis of Marmite preference.

## Methods

### Subject Recruitment

We enrolled 261 healthy adults who were over the age of 18 years, regardless of gender or race/ethnic group, to participate in this study. Study subjects were recruited from the office buildings in proximity of where DNAFit Ltd is located in London, UK. Recruitment was done by posting flyers for the study in public spaces and emailing further information to interested persons who contacted the study staff. Exclusion criteria for enrollment in the study included: 1) all employees of the companies involved in the study, 2) pregnant individuals, and 3) individuals with a food intolerance to Marmite and/or food allergies to any ingredients in Marmite. Additionally, subjects were screened the day of the study to exclude individuals that might have had a compromised sense of taste and/or smell at the time of data collection. All study subjects were provided the Participant Info Sheet in advance of the study, participated in an Informed Consent session, and given the opportunity to answer any questions that they had. All study subjects provided signed InformedConsent.

### Data Collection

After subjects completed enrollment in the study we collected phenotype data via a short questionnaire including 42 items. Questionnaire design followed standard survey methodology, generally accepted practices for sensitive questions,and specific guidelines from the UK ONS and UK Census for the study population. We collected demographic information (age, gender, nationality, ethnic group), phenotype information in response to tasting interventions (taste perception), additional phenotype information (food preference, food frequency), and information for use as covariates in the genetic association analysis (smoking status, and “supertaster” status).

Study subjects participated in two tasting interventions: 1) tasting ½ serving of Marmite (~2g) on their tongue for ~10sec, and 2) tasting a filter paper strip containing phenylthiocarbamide (PTC) on their tongue for ~10sec. PTC is commonly used to test for bitter tasting ability and has been validated as a method to report on the TAS2R38 genetic haplotypes. After each tasting, subjects were asked to describe the taste using a series of yes/no questions and then rate how much they enjoyed the taste on a Likert scale. The order of the two tasting interventions was randomized among subjects using an A/B scheme.

Study subjects provided a non-invasive biological sample for genotyping. Oragene OCD-100 cheek swab saliva kits were used for sample collection. Sample processing, DNA extraction, hybridization, and genotyping were done using standard Illumina protocols and equipment at an accredited UK HTA genomics facility. Illumina GSA bead-chip arrays with ~650,000 SNPs were used for genotyping. Saliva samples, DNA, and genetic information existed only in de-identified formatthroughout the study.

### Phenotype Data Analysis

Questionnaire data was collected in paper format, manually entered into a secure database by the PI, and then spot-checked for errors by a separate member of the study staff. Data entry, validation, and cleaning was performed in de-identified format. Summary statistics from questionnaire data were generated in Excel. Phenotype data was analyzed by standard statistical correlation methods using Excel and R software.

### Genetic Association Studies

Genetic association studies were performed to identify SNPs associated with Marmite taste preference. Three levels of analysis were performed: 1) a genome-wide association study (GWAS) using a total of 642,824 SNPs; 2) a general candidate gene approach including ~250 genes known to be involved in taste/olfactory processes; and 3) a specific candidate gene approach focusing on the TAS2R38 gene involved in bitter taste perception.

We tested 2 phenotypes related to Marmite taste preference: 1) status as a Marmite “lover” vs “hater”, and 2) rating of the taste of Marmite. We tested 2 phenotypes related to bitter tasting ability: 1) whether PTC tasted bitter, and 2) rating of the taste of PTC. When using a case/control model, cases were identified through a combination of subjects questionnaire responses to the Marmite taste intervention and self-report of being a Marmite “hater” or “lover”. In cases where the subjects result of the direct Marmite taste test conflicted with the subjects self-report of Marmite taste preference, the result of the Marmite taste test from this study was used to assign case/control status. When using a continuous model, individuals were classified as Marmite lovers, haters, or neutral; as well as liking, disliking, or neutral preference to the taste of Marmite.

We performed genome-wide association studies (GWAS) in Plink using: 1) basic association by chi-square and Fishers exact tests, and 2) a linear/logistic regression model with known covariates. All GWAS analyses followed standard practices, including: exclusion of low quality genotype samples, filtering of SNPs with low MAF, filtering of SNPs that deviate from HWE, evaluation of Q-Q plots, correction for any genomic inflation (lambda), and careful consideration of ethnic stratification. We set our statistical thresholds as: genome-wide significance (p<5x10^−8^), Bonferroni corrected significance (p<7.78x10-8), and nominal significance (p<1x105). We set our effect size estimates as: small effect (OR=1.0-1.5), medium effect ( 0R=1.5-9.0), and large effect (OR>9.0). Regression modeling was performed using the following covariates: 1) gender [binary], 2) age [continuous], and 3) smoking status [categorical]. All analyses were performed on the entire cohort (n=261) as well as the subgroup of white Europeans (n=213). We performed separate stratified analyses for men and women.

We performed candidate gene analyses in Plink and in Excel for: 1) a set of 252 genes involved in taste/olfactory processes, 2) 3 individual SNPs in the TAS2R38 gene, and 3) 2 haplotypes of the SNPs in the TAS2R38 gene. All candidate gene analyses followed standard practices, and included the same methodological considerations as described for the GWAS. Statistical analysis was performed using chi-square and Fishers exact tests to identify whether candidate genes and/or candidate SNPs reached nominal significance (p<0.05).

### Ethics Statement

This study has been reviewed and given favourable opinion by the Western Institutional Review Board research ethics committee (WIRB^®^ Protocol #20170967, Approval Date May 5th 2017). This study has been conducted using the highest level of ethics guidelines (ie. UK NHS guidelines, GCP, HIPPA). This study has been conducted in accordance with all legal requirements under the UK Human Tissue Act of 2004 and the UK Data Protection Act of 1998. All subjects provided signed Informed Consent.

